# Loss of Myeloid Cell-Specific β2-Adrenergic Receptor Expression Ameliorates Cardiac Function and Remodeling after Acute Ischemia

**DOI:** 10.1101/2023.11.27.568873

**Authors:** Tapas K. Nayak, Anamika Bajpai, Viren Patwa, Rhonda L. Carter, Nitya Enjamuri, Erhe Gao, Yang K. Xiang, Douglas G. Tilley

## Abstract

Myeloid cells, including neutrophils, monocytes and macrophages, accumulate quickly after ischemic injury in the heart where they play integral roles in the regulation of inflammation and repair. We previously reported that deletion of β2-adrenergic receptor (β2AR) in all cells of hematopoietic origin resulted in generalized disruption of immune cell responsiveness to injury, but with unknown impact on myeloid cells specifically. To investigate this, we crossed floxed β2AR (F/F) mice with myeloid cell-expressing Cre (LysM-Cre) mice to generate myeloid cell-specific β2AR knockout mice (LB2) and subjected them to myocardial infarction (MI). Via echocardiography and immunohistochemical analyses, LB2 mice displayed better cardiac function and less fibrotic remodeling after MI than the control lines. Despite similar accumulation of myeloid cell subsets in the heart at 1-day post-MI, LB2 mice displayed reduced numbers of Nu by 4 days post-MI, suggesting LB2 hearts have enhanced capacity for Nu efferocytosis. Indeed, bone marrow-derived macrophage (BMDM)-mediated efferocytosis of Nu was enhanced in LB2-versus F/F-derived cells in vitro. Mechanistically, several pro-efferocytosis-related genes were increased in LB2 myeloid cells, with annexin A1 (*Anxa1*) in particular elevated in several myeloid cell types following MI. Accordingly, shRNA-mediated knockdown of *Anxa1* in LB2 bone marrow prior to transplantation into irradiated LB2 mice reduced Mac-induced Nu efferocytosis in vitro and prevented the ameliorative effects of myeloid cell-specific β2AR deletion on cardiac function and fibrosis following MI in vivo. Altogether, our data reveal a previously unrecognized role for β2AR in the regulation of myeloid cell-dependent efferocytosis in the heart following injury.

## Introduction

Acute ischemic injury, such as myocardial infarction (MI), is a major contributor of cardiovascular disease, a leading cause of mortality in the U.S., which affects millions of patients and costs billions of dollars annually^1, 2^. Myeloid cells, including neutrophils (Nu), monocytes (Mon) and macrophages (Mac), normally constitute a small fraction of the adult heart but play major roles in the response to acute ischemic injury, for instance scavenging dead cells and secreting numerous factors such as cytokines and chemokines to promote the accumulation of additional inflammatory cells^3–9^ . Myeloid cells perform several functions including efferocytosis (receptor-mediated phagocytosis), the clearing of dead cells and debris from a site of injury^10, 11^. Nu clearance by Mac from the infarct site in the heart is critical for inflammation resolution and scar formation since the presence of dead/dying aged Nu triggers secondary inflammation that can lead to further damage and tissue necrosis^12^. Mac-dependent efferocytosis is one of the dominant mechanisms of aged Nu clearance and can be mediated by several receptor-ligand interactions. Recent work has detailed the importance of efferocytosis following cardiac injury and some of the key molecular players involved, including annexin A1 (Anxa1)^9, 13–15^ . Indeed, several studies have demonstrated that myeloid cell (Nu or Mac)-produced Anxa1 acts to enhance Nu apoptosis and subsequent Mac-mediated efferocytosis to resolve inflammation^4, 16–21^. Ultimately, more efficient clearance of apoptotic Nu/debris leads to better cardiac function and remodeling outcomes.

As reports accumulate each year better defining myeloid cell subpopulations and their plasticity within the heart, either normally or following injury, their potential for modulation toward promoting improved cardiac function and remodeling outcomes continues to grow. Myeloid cells express an extensive repertoire of cell surface receptors that regulate their function and responsiveness to injury, including G protein-coupled receptors (GPCRs), stimulation of which can induce rapid alterations in cell polarity, chemotaxis, adhesion and migration in myeloid cells^22–24^. Chemokine GPCRs in particular have been a major focus as regulators of immune cells, however many other receptor types, including neurohormone GPCRs, are expressed on myeloid cells and can impact their function^25^. Indeed, the β2-adrenergic receptor (β2AR) is expressed on numerous immune cell types and has been shown to regulate an array of functions, including inflammation, egress, migration and immunosuppression^26–30^.

We previously reported that deletion of β2AR in all cells of hematopoietic origin in mice significantly reduced immune cell recruitment to the heart following experimental myocardial infarction (MI), resulting in left ventricular rupture and 100% mortality within the first 2 weeks post-MI^31, 32^. Unclear from these studies was the impact on cardiac remodeling outcomes of β2AR deletion specifically in myeloid cells since they represent the first responders to acute cardiac injury. Thus, to investigate the role of myeloid cell-specific β2AR in the response to acute cardiac injury, we crossed floxed β2AR and constitutive myeloid cell-expressing Cre (LysM-Cre) mice to generate myeloid cell-specific β2AR knockout mice. Here we demonstrate that these mice are unexpectedly protected against maladaptive cardiac remodeling post-MI, an effect mediated through enhanced Nu efferocytosis. Additionally, we show that the increased efferocytosis observed in β2AR-deficient myeloid cells is Anxa1-dependent and that by decreasing Anxa1 expression in β2AR knockout bone marrow cells, normal in vitro efferocytosis and in vivo cardiac remodeling responses are restored. Thus, myeloid cell-specific β2AR deletion improves post-ischemic injury-induced cardiac function and fibrotic remodeling via enhanced efferocytosis-mediated Nu clearance.

## Materials and Methods

### Animal Studies

Experiments using mice were conducted under the National Institutes of Health (NIH) Guide for the Care and Use of Laboratory Animals, and also approved by the Institutional Animal Care and Use Committee (IACUC) at Temple University with Animal Protocols 4891 and 4902. β2AR floxed mice (FL/FL)^33^ were a gift from Dr Gerald Karsenty (Columbia University) and were used for crossbreeding with LysMcre (LMC) (#004781; The Jackson Laboratory, ME, USA) to develop myeloid cell-specific β2AR knockout mice (LB2). All mice were housed with ad libitum food and water and genotyped using flox site- and Cre-specific primers before experimental procedures (Supplemental Table 1).

### Coronary Artery Occlusion Surgery

Both male and female mice (12-14 weeks old) were used for experimental myocardial infarction (MI) as reported previously^31, 32, 34^. Briefly, mice were anesthetized with 3% isoflurane inhalation, after which a small skin incision was made over the left chest, and a purse suture was made, allowing the pectoral muscles to be dissected and retracted. The 4th intercostal space was then exposed, and a small hole was made to gently extract the heart. Once the left main descending coronary artery (LCA) was located, it was sutured and ligated approximately 3 mm from its origin with a 6-0 silk suture, after which the heart was placed back into the intra-thoracic space and the chest was closed. Sham-operated mice were treated similarly, except the LCA was not occluded. Immediately after surgery, all mice received a single subcutaneous dose of buprenorphine at 0.3 mg/kg for pain alleviation.

### Echocardiography

Cardiac function was assessed by transthoracic 2-dimentional M-mode (short axis) echocardiography using the VisualSonic Vevo 2100 Imaging System (FUJIFILM, Toronto, ON). Prior to echocardiography, chest fur was removed by Nair for clear image acquisition. Mice were anesthetized by 3% isoflurane inhalation and maintained between 1.0-1.5% throughout the imaging procedure. Echocardiography LV-trace data analysis was performed by using Vevo Lab 5.5.0 software.

### Lentivirus Transduction and Bone Marrow Transplant (BMT)

Bone marrow cells from the femurs of LB2 mice were isolated and transduced with either scrambled or Annexin A1 (Anxa1) shRNA using TR30030 pRFP-C-shLenti vector (Origene Technologies Inc. MD, USA) at an MOI of 150. Lentivirus transduction was performed in serum free RPMI-1460 (w/o antibiotic and antimycotic) in the presence of ViralEntry™ Transduction Enhancer (Applied Biological Materials Inc. ABM, BC, Canada). A non-effective shRNA cassette in the TR30030 pRFP-C-shLenti vector plasmid was used as a scrambled control. After 2h, cells were washed in 1x PBS and injected immediately to irradiated mice. 8-week-old LB2 male mice were irradiated using RS-2000 X-ray irradiator with 950 rads to deplete recipient bone marrow cells. Sex- and age-matched donor LB2 mouse bone marrow cells infected with lentivirus (as described above) were adoptively transferred (10x10^6^ live cells/mouse) into isoflurane-anesthetized mice within 24h of irradiation via retroorbital injection. Immediately after the procedure, all BMT recipient mice received a single sub-Q dose of buprenorphine at 0.3 mg/kg to alleviate pain. All the animals were allowed to reconstitute the bone marrow for 1-month prior experiments. Upon the termination of the experiments, the success of BM cell reconstitution was confirmed for each mouse by RT-qPCR.

### Heart Perfusion and Digestion

Mice were euthanized by isoflurane overdose, followed by cervical dislocation as per Temple university IACUC approval. The diaphragm was surgically exposed without damaging any organs and perfused with excess 1X PBS (cold) to reduce vascular immune cell contamination. The heart was harvested and stored in RMPI-1460 on ice until tissue processing. Using an industrial blade, the heart was minced ∼1mm^3^ in size (and until a smooth consistency was observed) and collected in a 50 mL falcon tube containing digestion buffer composed of Collagenase type-2 (450 U/mL), Hyaluronidase (60 U/mL) and DNase I (60 U/mL) in serum-free RPMI-1460. The digestion was carried out in an incubator set at 37°C with gentle horizontal shanking for 1 h. FBS was added to a final concentration of 10% (v/v) and the digested tissues were passed through 40 µM cell strainer. The homogenized samples were centrifuged at 400 x g for 5 min at 4°C and cell pellets were subjected to 1X ACK lysing buffer for 2 min to remove RBC contamination. Once RBC lysis is complete, 2 volumes of 1X PBS was added and centrifuged as above. The resultant cells were resuspended in complete RPMI-1460 and stored at 4°C until downstream application.

### Cell Culture and Primary Cell Isolation

L929 cells were maintained in complete Dulbecco′s Modified Eagle′s Medium (DMEM) supplemented with penicillin, streptomycin, and amphotericin-B (PSF) and 10% Fetal bovine serum (FBS) at 37°C under a humidified incubator with 5% CO2. Cell culture supernatants containing M-CSF were collected twice a week, passed through 0.22 µM sterile filter and stored at -20°C until use for macrophage differentiation.

Mouse bone marrow cells were isolated as described by others^35, 36^. Briefly, both femurs and tibias were flushed with RPMI-1460 supplemented with 2mM EDTA and cells were collected in a 50 mL falcon tube. Cells were centrifuged at 300 x g for 5 min at room temperature (RT). Red blood cells (RBC) were lysed using 1x Ammonium-Chloride-Potassium (ACK) lysing buffer followed by filtered through 40 µM cell strainer and were spun down as above. The resultant cell pellets were resuspended in respective media as per downstream applications. For bone marrow-derived macrophage (BMDM) culture, cells were resuspended in complete RPMI-1460 containing 20% L929 conditioned media at a seeding density of 10x10^6^ cells/100 mm dish. After 4 days, media was replaced, and cells were cultured for another 2 days. On day 6, BMDM were detached by 2mM EDTA treatment and re-seeded in tissue culture plates as per downstream experiments.

### Cell Enrichment by Magnetic Beads

CD11b+ cells (130-049-601) and BM neutrophils (130-097-658) were isolated using Miltenyi Biotec (Bergisch Gladbach, Germany) microbeads as per manufacturer’s instructions. For CD11b+ cell isolation from digested hearts, cell pellets were precleared with debris removal solution (130-109-398, Miltenyi Biotec). Briefly, cells were spun at 300 x g for 10 min at 4°C and supernatants were aspirated completely. Cells were resuspended with an appropriate volume of debris removal solution in a 15 mL tube, overlaid by 4mL of cold 1x PBS and spun as above for phase separation in a swinging bucket centrifuge with full acceleration and full brake. The top two phases containing buffer and interphase were discarded and the tube was filled to a final volume of 15 mL using cold 1X PBS. The solution was mixed gently by inverting three times and spun. Cells were resuspended in freshly prepared isolation buffer (1X PBS, pH 7.2) supplemented with 2 mM EDTA, 0.5% FBS (v/v) and kept on ice until enrichment. After determining cell number, cells were incubated with CD11b microbeads (20 uL per 10 x 10^6^ cells), which was diluted in isolation buffer (20 uL per 10 x 10^6^ cells). After 15 min incubation on ice, cells were washed and resuspended in isolation buffer (500uL/10^8^ cells) for column-based magnetic separation. For neutrophil enrichment, 50 x 10^6^ BM cells were resuspended in chilled isolation buffer (200 uL) containing 20 uL neutrophil biotin antibody cocktail. After 15 min, cells were washed and incubated with anti-biotin microbeads (100 uL) diluted in isolation buffer (400 uL) at 4°C. After 15 min incubation, cells were washed and resuspended in isolation buffer (500uL/10^8^ cells) for column based magnetic separation. Microbead-labelled cells were passed through Miltenyi Biotec MS columns (130-122-727) placed on a MiniMACS™ Starting Kit (130-090-321), and cell fractions were collected as per the cell-type enrichment (CD11b+ cells; negative selection and neutrophils; negative selection).

### Flow Cytometry

Equal volume of single cell suspension from digested hearts were taken in 5 mL polystyrene round-bottom Falcon tubes and spun at 1000 rpm for 5 min at 4 °C. Then cells were incubated in fluorophore labeled antibody (as described in Supplemental Table 2) diluted in FACS buffer (1X PBS, 10% FBS). After 45 min on ice, cells were washed in FACS buffer. For intracellular staining, BD cytofix/cytoperm fixation/permeabilization solution kit was used as per manufacturer’s protocol. Fluorophore-matched IgGs were used as isotype controls. The FcR blocking reagent (Miltenyi Biotec, Germany) was used 15 min prior to the primary antibody incubation in order to block the non-specific binding of antibodies to Fc receptors. Cells were acquired in BD LSR-II flow cytometer using FACS diva software and FCS files were analyzed by the Flowjo v10.5.0 software (BD Biosciences, CA, USA). For absolute counting, total cells were normalized to mg tissue weight. For ex vivo experiments, 1 x 10^6^ cells were taken for staining and a total of 1 x 10^5^ cells were acquired per sample. UltraComp eBeads™ (Thermo Scientific, USA) was used for fluorophore channel compensation. High FSC/SSC events were gated, followed by the exclusion of doublets (FSC-A vs FSC-H). CD45+ cells were selected for further analysis of CD11b+ Ly6G+ neutrophils, CD11b+ Ly6Chi/lo monocytes, CD11b+ F4/80+ macrophages.

### Reverse Transcription Quantitative Polymerase Chain Reaction (RT-qPCR)

Total RNA was isolated, and cDNA was synthesized using PureLink™ RNA Mini Kit and High-Capacity cDNA Reverse Transcription Kit (Thermo Fisher scientific, USA) respectively. RT-qPCR was carried out with equal amount of cDNA by PowerUp™ SYBR™ Green Master Mix (Thermo Fisher scientific, USA) in triplicate using specific primers (Supplemental Table 1). All the primers were designed using Primer3Plus website (https://www.primer3plus.com). RT-qPCR data was analyzed using Quant studio design and analysis software v1.5.1 by comparative CT method (ΔΔCT) and normalized to glyceraldehyde 3-phosphate dehydrogenase (GAPDH) gene expression.

### Efferocytosis Assay

Freshly isolated neutrophils were stained with CellTracker™ Green 5-chloromethylfluorescein diacetate (CMFDA) dye (Thermo Fisher scientific, USA) at 5µM in serum free RPMI-1460 (SFM) for 30 minutes in the incubator, followed by washing in SFM. Both BMDM and CMFDA-stained neutrophils were treated with isoproterenol at 10µM for 16h. The efferocytosis assay was carried out by co-culturing BMDM-neutrophils at a ratio of 1:3 as per previous report (PMID_32336197) for 45 min. Cells were washed twice in 1X PBS to remove unbound neutrophils and BMDM were detached using enzyme-free cell dissociation buffer (Thermo Fisher scientific, USA). Then cells were processed for flow cytometry-based analysis as described above after labelling with F4/80 antibody.

### Tissue Processing and Masson’s Trichrome Staining

After euthanasia as described above, excised hearts were collected and perfused with 1X PBS containing 1 M potassium chloride on ice for diastolic heart arrest. Hearts were fixed in an excess volume of 4% paraformaldehyde, paraffin embedded and sectioned at a thickness of 5 µM. Deparaffinized and rehydrated sections were processed for Masson’s trichrome staining (Sigma-Aldrich, USA). Sections were visualized on a Nikon Eclipse microscope at 20X magnification. Infarct sizes were calculated using NIS elements software and interstitial fibrosis was calculated using ImageJ software.

### Statistical Analysis

GraphPad prism 9 (GraphPad Software Inc. USA) was used for statistical analysis. Data were represented Mean ± SEM. Comparison between different groups were carried out by either one-way or two-way ANOVA (repeated measures or mixed model) with Tukey’s multiple comparisons test. Unpaired student’s t-test was used for comparison between mean of two variables. *p* < 0.05 was considered statistically significant between groups.

## Results

### Mice with myeloid cell-specific β2AR deletion display improved cardiac function and remodeling post-MI

To investigate the role of myeloid cell-specific β2AR in the response to acute cardiac injury, we crossed floxed β2AR (FL/FL) and constitutive myeloid cell-expressing Cre (LysM-Cre) mice to generate myeloid cell-specific β2AR knockout mice (LB2, Fig. 1A). Via RT-qPCR, a significant reduction in β2AR expression was observed in total bone marrow (BM) cells from LB2 versus FL/FL and LMC BM cells (Fig. 1B), while almost complete ablation of β2AR was observed in LB2 versus FL/FL and LMC bone marrow-derived macrophages (BMDM) (Fig. 1C). To assess the impact of myeloid cell-specific β2AR deletion on cardiac function and remodeling following injury, we subjected FL/FL, LMC and LB2 mice to sham or MI surgery and monitored cardiac function via echocardiography over time (Fig. 1D). M-mode echocardiography revealed a significant preservation of several cardiac parameters in LB2 mice versus the control groups at 3 days post-MI (Fig. 1E, Supplemental Fig. 1), including % ejection fraction (EF, Fig. 1F) and % fractional shortening (FS, Fig. 1G). By 14 days post-MI, %EF and %FS were still significantly elevated in LB2 mice compared to the control groups.

**Figure 1.**
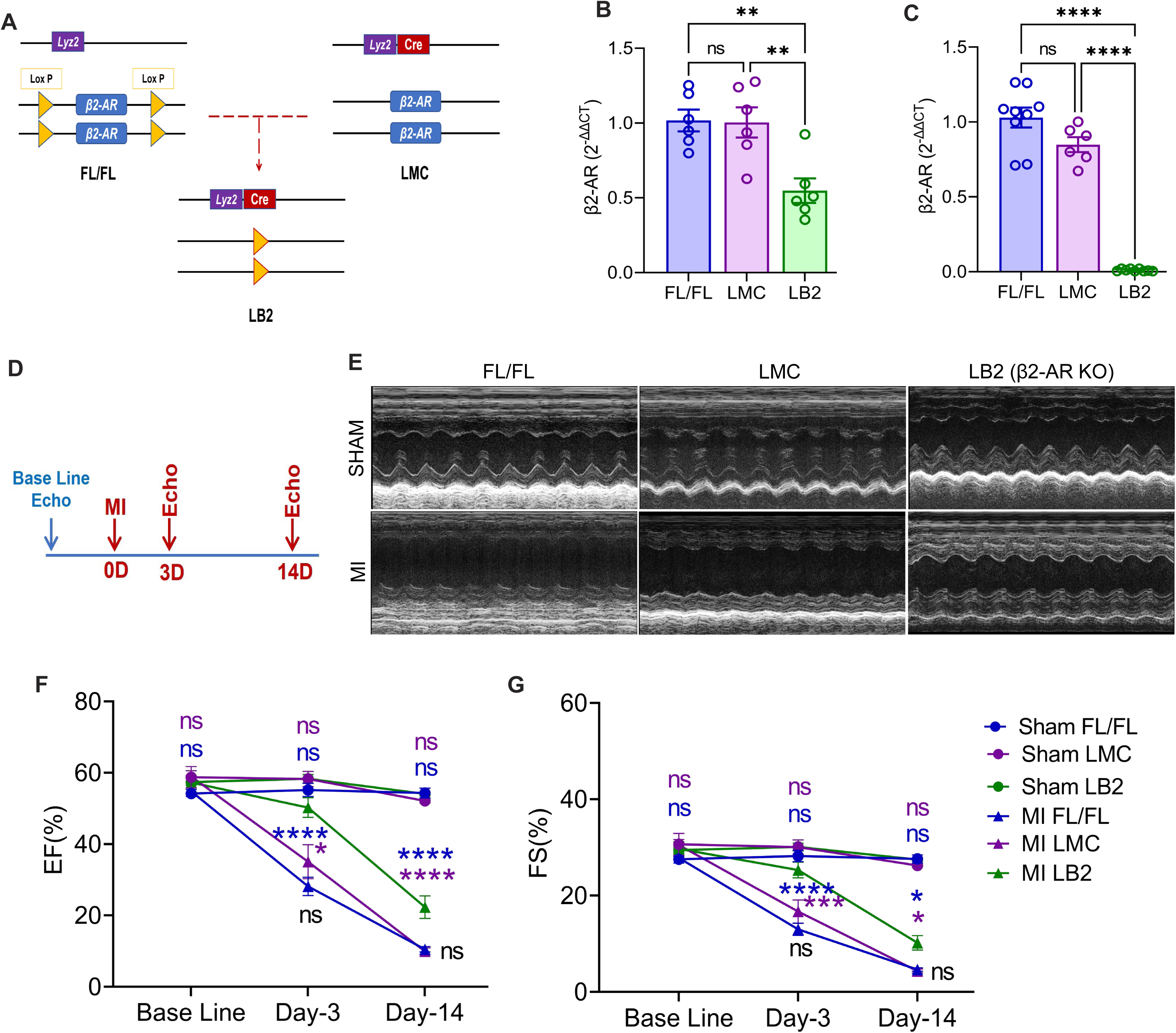
Mice with myeloid cell-specific β2AR deletion display improved post-MI cardiac function. (A) Graphical illustration showing myeloid cell-specific β2AR deletion achieved via crossing β2AR^flox/flox^ (FL/FL) and LysMCre (LMC) mice, resulting in myeloid cell-specific β2AR KO mice (LB2). Histograms depicting *β2AR* expression as detected via RT-qPCR in bone marrow cells (B, n=6/group) and differentiated bone marrow-derived macrophages (C, n=8 for FL/FL, n=6 for LMC, n=9 for LB2). (D) Study design showing timeline of surgery and echocardiography measurements. (E) Representative echocardiography images (M-mode short axis) of sham (upper panels) or MI (lower panels) FL/FL, LMC or LB2 mice. Histogram depicting percent ejection fraction (%EF, F) and fractional shortening (%FS, G) at baseline (BL), Day-3 and Day-14 post-MI for sham or MI FL/FL (blue), LMC (purple) or LB2 (green) mice Data are Mean ± SEM of independent experiments. ns, non-significant, *p < 0.05; **p < 0.01; ****p < 0.0001, One way ANOVA with Tukey’s post-hoc test (B, C and D) or Two-way ANOVA (mixed model) with Tukey’s post-hoc test (F, G). n=11 for sham FL/FL, n=9 for sham LMC, n=12 for LB2, n=10 for MI FL/FL, n=5 for MI LMC, n=11 for MI LB2 (day 3), n=12 for MI FL/FL, n=9 for MI LMC, n=6 for MI LB2 (day 14).

Since cardiac function was improved in the LB2 mice versus the control lines, we surmised that cardiac remodeling events may also be ameliorated. Via echocardiography the LB2 mice displayed better preservation of wall thicknesses post-MI, more like the sham-operated groups, than FL/FL and LMC MI-operated mice (Supplemental Fig. 1), suggesting that along with preserved function myeloid cell-specific β2AR deletion results in less pathologic wall thinning. Indeed, via Masson’s trichrome (MT) staining at 14 days post-MI, we observed a reduction in % infarct lengths and LV wall thinning in LB2 mice as compared to either control line (Fig. 2A, B). Additionally, interstitial fibrosis in the border zone of LB2 mice was significantly reduced versus the control lines (Fig. 2C, D), while the remote zone was devoid of fibrosis in all lines. Notably, FL/FL and LMC control mice did not differ in any parameter assessed, thus FL/FL mice were subsequently used as controls for the remainder of the study to reduce duplicative animal usage. Collectively, these data suggest that the deletion of myeloid cell-specific β2AR enacts early benefits against ischemic cardiac injury to better sustain cardiac function and decrease infarct expansion and fibrosis over time.

**Figure 2.**
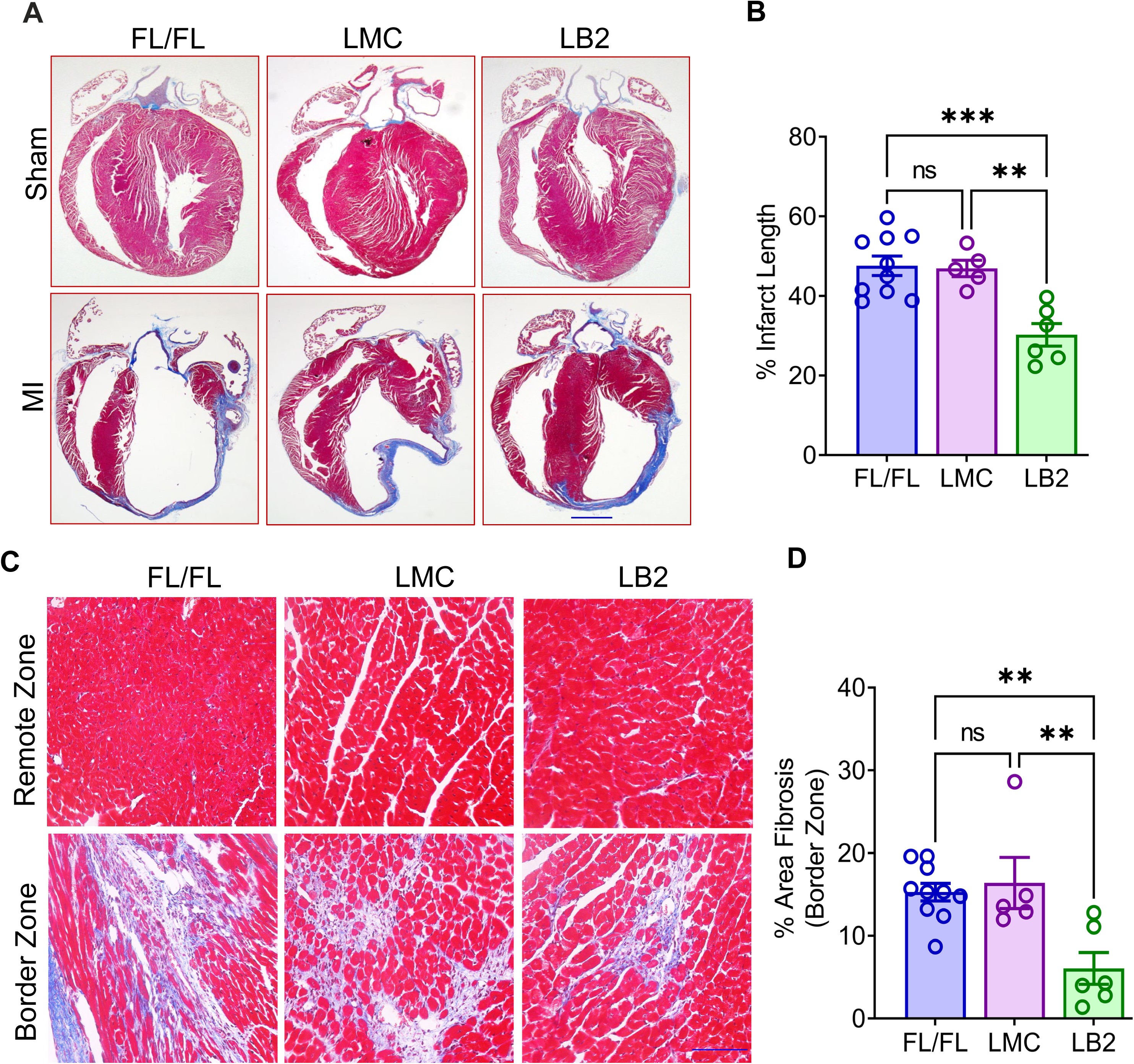
Mice with myeloid cell-specific β2AR deletion have reduced infarct length and cardiac fibrosis post-MI. (A) Masson’s Trichome (MT) staining for sham and 14-Day post-MI hearts of FL/FL, LMC and LB2 mice, scale bar = 1000 µm. (B) Graph depicting % infarct length at 14 days post-MI. (C) Bright field microscopic images (20X, scale bar = 100 µm) of MT staining in the remote zone (upper) and border zone (lower) with quantification (D) of % area fibrosis in the border zone. Data are Mean ± SEM of independent experiments. n=10 for FL/FL, n=5 for LMC, n=6 for LB2. ns, non-significant, **p < 0.01; ***p < 0.001, One way ANOVA with Tukey’s post-hoc test (B, D).

### Myeloid cell-specific deletion of β2AR enhances post-MI cardiac neutrophil clearance

Although myeloid cells, primarily Nu, Mon and Mac, only constitute about 7% of the heart basally, they play important roles in regulating cardiac structure and function in a spatiotemporal manner following injury^3, 4, 8, 37–39^. Since LB2 mice exhibited better cardiac function post-MI, we sought to assess whether β2AR deletion impacts the recruitment or retention of the major myeloid cell populations in the heart. Thus, single cell suspensions from enzymatically digested post-MI hearts were analyzed for quantification of phenotypically different myeloid cells using specific markers and gating strategies (Fig. 3A, B). All CD45+ immune cells were initially gated, then Nu, Mon and Mac were quantified using CD11b+ Ly6G+, CD11b+Ly6Chi/lo and CD11b+F4/80+ gates, respectively. At 1-day post-MI, the number of each myeloid cell population recruited to the heart did not significantly differ between LB2 and FL/FL mice (Fig. 3C). However, by 4 days post-MI, CD45+CD11b+Ly6Clo Mon and CD45+CD11b+Ly6G+ Nu were significantly reduced in LB2 versus FL/FL mouse hearts (Fig. 3D). Unlike Ly6Clo Mon, mature Nu are short-lived and mainly divide in the bone marrow and to a lesser extent in the spleen^40^. Considering this, and that the initial recruitment of Ly6Clo Mon and Nu was not different in LB2 hearts at 1-day post-MI, our data suggest that the decreased amount of these cells in LB2 hearts at 4 days post-MI may be due to enhanced clearance.

**Figure 3.**
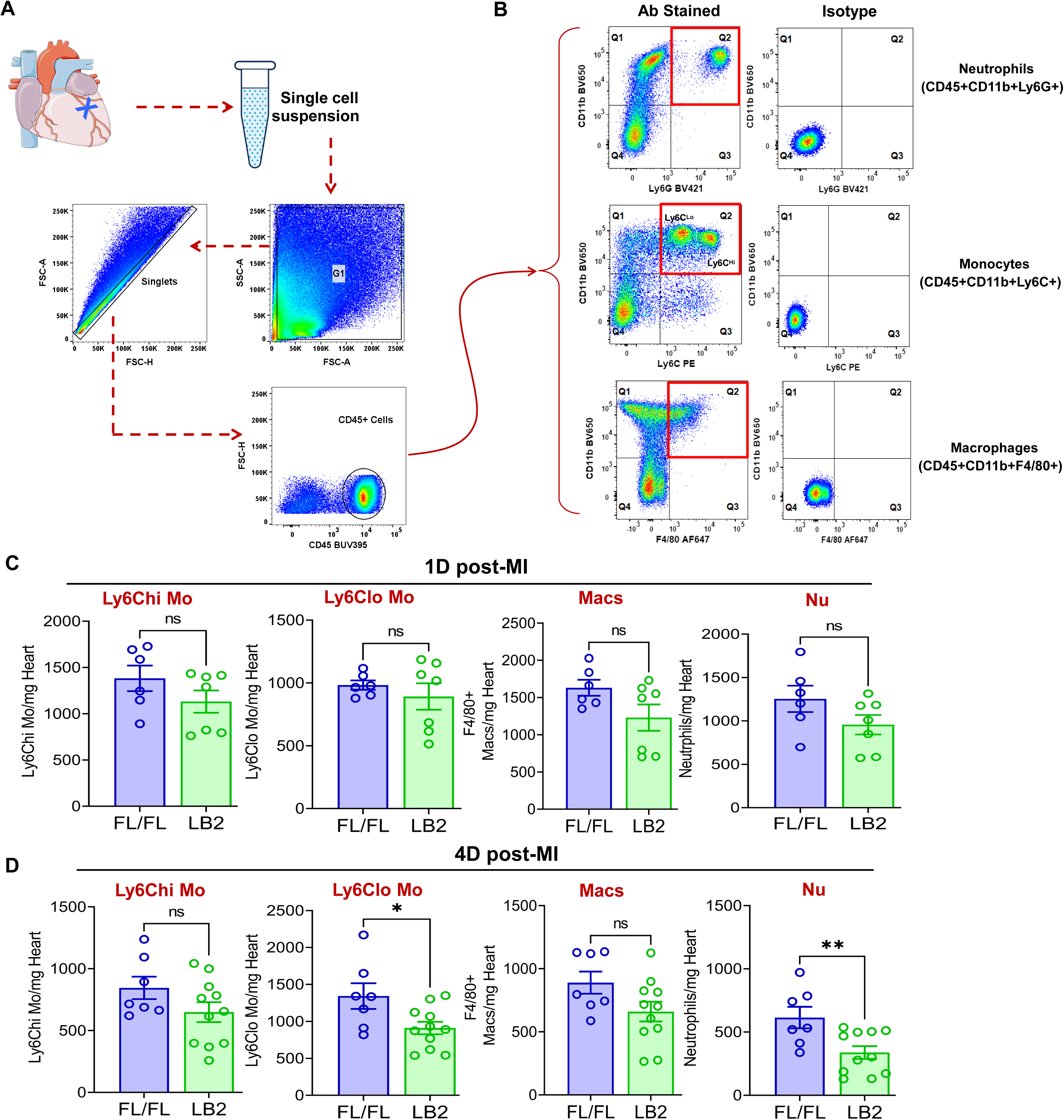
Myeloid cell-specific deletion of β2AR reduces post-MI cardiac neutrophil clearance. (A) Flow chart representation of gating strategies used for flow cytometry-based immune cell profiling. (B) Flowjo analysis of representative pseudo-colored dot plots of CD45+ gated antibody-stained (left column) Nu (top), Mon (middle) and Mac (bottom) along with corresponding isotype controls (right column). Histograms representing absolute number of immune cell types indicated, CD11b+Ly6Chi Mon, CD11b+Ly6Clo Mon, CD11b+F4/80+ Mac and CD11b+Ly6G+ Nu, normalized per mg of heart weight at 1-Day (C) and 4-Day (D) post-MI. Data are Mean ± SEM of independent experiments. n=6 (1-day) or 7 (4-day) for FL/FL, n=7 (1-day) or 11 (4-day) for LB2. ns, non-significant, *p < 0.05; **p < 0.01, Unpaired student’s t-test.

### Annexin A1 expression is increased in β2AR-deficient myeloid cells post-MI

Following cardiac injury, Nu quickly migrate to the infarct area where they become activated and play a dominant role in the acute inflammatory response and early engulfment of cellular debris^41, 42^. Due to their short life span, Nu begin to die in the days following MI and are phagocytosed by Mac via a variety of receptor-ligand mediated interactions in a process called efferocytosis, which limits further Nu-mediated inflammation^10, 11^. Accordingly, inefficient or prolonged Nu clearance further promotes sustained secondary tissue death and inflammation^11, 12, 43^. Since we observed a greater reduction in the number of Nu in LB2 hearts at 4 days post-MI, despite similar levels of their recruitment at 1 day-post-MI, we hypothesized that myeloid cell-specific β2AR deletion promotes more efficient Nu efferocytosis after infarction. To test this, BM cells were screened for the expression of different efferocytosis-related genes by RT-qPCR. We observed that *Anxa1*, *Mertk*, *Axl*, *Stab1* and *Stab2*, genes with known roles in efferocytosis, were significantly altered in LB2 versus FL/FL BM cells at baseline (Fig. 4A). Further, we observed significantly higher expression of *Anxa1* in both BM cells and enriched cardiac CD11b+ myeloid cells isolated from LB2 versus FL/FL mice at 1-day post-MI (Fig. 4B-F). Flow cytometry analysis was carried out at 4 days post-MI to determine which immune cell subtypes displayed enhanced Anxa1 expression in LB2 mice. Via median fluorescence intensity (MFI) analysis we detected significantly more AnxA1 protein expression in LB2 versus FL/FL cardiac CD11b+Ly6G+ Nu, CD11b+Ly6Chi Mon and CD11b+F4/80+ Mac, but not in CD11b+Ly6Clo monocytes (Fig. 4G-J). These data suggest that myeloid cell-specific β2AR deletion may facilitate Nu clearance after cardiac injury via increased AnxA1 expression.

**Figure 4.**
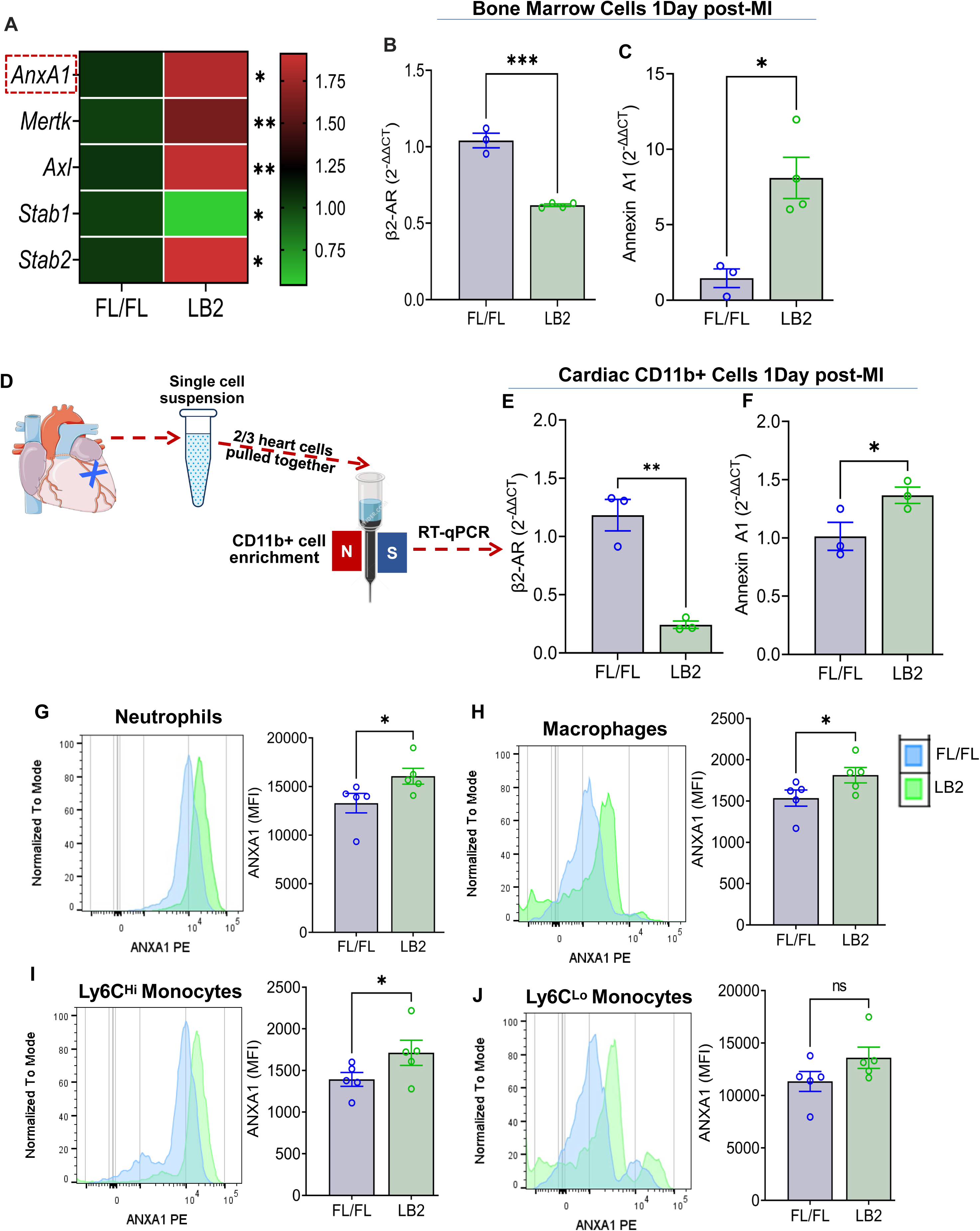
Annexin A1 expression is increased in β2AR-deficient myeloid cells post-MI. (A) RT-qPCR screening of various efferocytosis pathway genes was performed on total RNA isolated from whole bone marrow (BM) cells from FL/FL and LB2 mice, with the heatmap representing the differential expression pattern of *Anxa1*, *Mertk*, *Axl*, *Stab1* and *Stab2* genes between FL/FL and LB2 BM cells. Gene expression analysis of *Adrb2* (β2AR, B) and *Anxa1* (Annexin xA1, C) of 1-Day post-MI total BM cells, n=3 for MI FL/FL and 4 for MI LB2. (D) Schematic of magnetic microbead column-based cardiac CD11b+ cell enrichment at 1-Day post-MI, after which gene expression of β2AR (E) and AnxA1 (F) was assessed via RT-qPCR, n=3/group. Flow cytometry-based histogram analysis showing comparative AnxA1 protein expression via mean fluorescence intensity (MFI) among various phenotypically distinct myeloid cell types: Nu (G), Mac (H), Ly6Chi Mon (I) and Ly6Clo Mon (J) in 4-Day post-MI FL/FL and LB2 hearts, n=5/group. Data are Mean ± SEM of independent experiments. ns, non-significant, *p < 0.05; **p < 0.01, *** p < 0.001, Unpaired student’s t-test.

### Myeloid cell-specific deletion of β2AR enhances neutrophil efferocytosis in vitro

To determine whether LB2 myeloid cells undergo more efficient efferocytosis, we employed an *in vitro* assay in which we freshly isolated BM cells to concentrate Nu, the purity of which were assessed by CD11b+Ly6G+ FACS staining and consistently found to be ∼90% (Supplemental Fig. 2), or to generate BMDM. To model *in vivo* conditions post-MI, differentiated BMDM and CMFDA-stained Nu were each treated with 10 µM isoproterenol (ISO) for 16 h, before co-culture within their respective genotype groups at a ratio of 1:3 (BMDM : Nu) for 45 min (Fig. 5A). Via flow cytometry, Nu efferocytosis by BMDM (i.e. F4/80+CMFDA+ events) was shown to be significantly higher in the LB2 versus FL/FL-derived cells (Fig. 5B-D). Moreover, the MFI signal of CMFDA within F4/80+ gated cells was also significantly elevated (Fig. 5E). Notably, *Anxa1* gene expression was found to be significantly higher in LB2-derived Nu (Fig. 5F), although a modest but non-significant increase was also observed in the BMDM *in vitro* (Fig. 5G).

**Figure 5.**
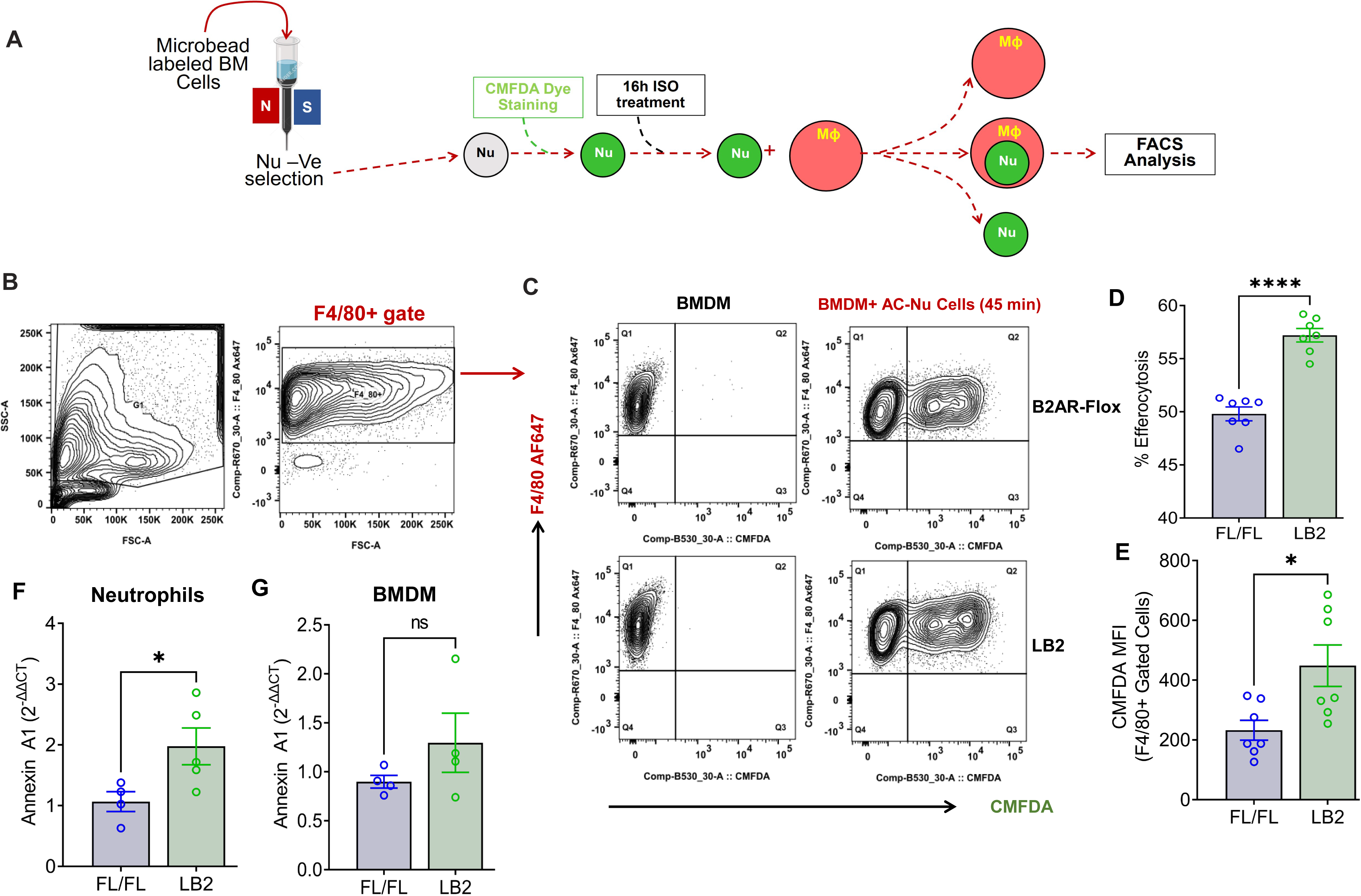
Myeloid cell-specific deletion of β2AR enhances neutrophil efferocytosis in vitro. (A) Schematic depicting experimental in vitro efferocytosis assay protocol in which Nu were freshly isolated from FL/FL or LB2 BM, loaded with CMFDA and treated with isoproterenol (ISO, 10µM) or vehicle (0.002% L-ascorbic acid) for 16 h prior to incubation with FL/FL or LB2 BMDM (also treated for 16 h with ISO) for 45 min at a 1:3 ratio. (B) FlowJo contour plot showing SSC-A/FSC-A and FSC-A/F4/80+ gate for Mac. (C) Contour plot of only Mac with (right column) or without (left column) CMFDA+ Nu. Quantification of Nu uptake efficiency into BMDM in terms of %efferocytosis (D) and F4/80+ gated CMFDA MFI (E), n=7 each. RT-qPCR based quantification of *Anxa1* gene expression in Nu (F) and BMDM (G) treated with ISO for 16 h, n=4 (F/F Nu), 5 (F/F BMDM), 5 (LB2 Nu), 4 (LB2 BMDM). Data are Mean ± SEM of independent experiments. ns, non-significant, *p < 0.05; **** p < 0.0001, Unpaired student’s t-test used.

### Lentivirus shRNA-mediated Anxa1 downregulation in LB2 neutrophils reduces their efferocytosis

Since AnxA1 is a pleotropic protein that modulates several immune functions including Nu apoptosis and efferocytosis by Mac^44–47^, we next tested whether downregulation of AnxA1 would restore LB2 myeloid cell efferocytosis efficiency to the level observed in FL/FL myeloid cells. We tested four different lentivirus clones (A-D, Supplemental Table 3) of mouse *Anxa1*-shRNA in BMDM for their gene silencing efficacy. RT-qPCR analysis showed all the clones were able to significantly decrease *Anxa1* expression in comparison to the scrambled shRNA control (Fig. 6A). Moreover, clones B and D showed maximum efficacy (∼50% reduction) as compared to clones A and C. Thus, clones B and D were mixed at a ratio of 1:1 and used in all the subsequent experiments. We have previously used lentivirus-mediated infection of bone marrow cells prior to transplantation as a method to manipulate immune cells responsiveness to cardiac injury in vivo^32^, thus using this approach we aimed to knockdown *Anxa1* in all cells of hematopoietic origin. For this, freshly isolated LB2 bone marrow cells were transduced with CTL or AnxA1 (Clones B/D) shRNA lentivirus prior to bone marrow transplantation (BMT) into lethally irradiated LB2 mice (Fig. 6B). The mice were subsequently used for experiments following at least 4 weeks of BM reconstitution, after which their BM was harvested and *Anxa1* expression confirmed via RT-qPCR (Fig. 6C). Initially, Nu were enriched from the BM and efferocytosis assays were carried out, as described in Fig. 5A, by incubating LB2 BMDM with either LB2 BMT CTL shRNA Nu or LB2 *Anxa1* shRNA Nu. As anticipated, efferocytosis of Nu isolated from LB2 BMT mice with *Anxa1* shRNA-mediated knockdown was found to be significantly reduced compared to Nu isolated from LB2 BMT mice with CTL shRNA transduction (Fig. 6D, E). These data support the notion that myeloid cell-specific β2AR deletion leads to enhanced Nu efferocytosis via increased expression of AnxA1.

**Figure 6.**
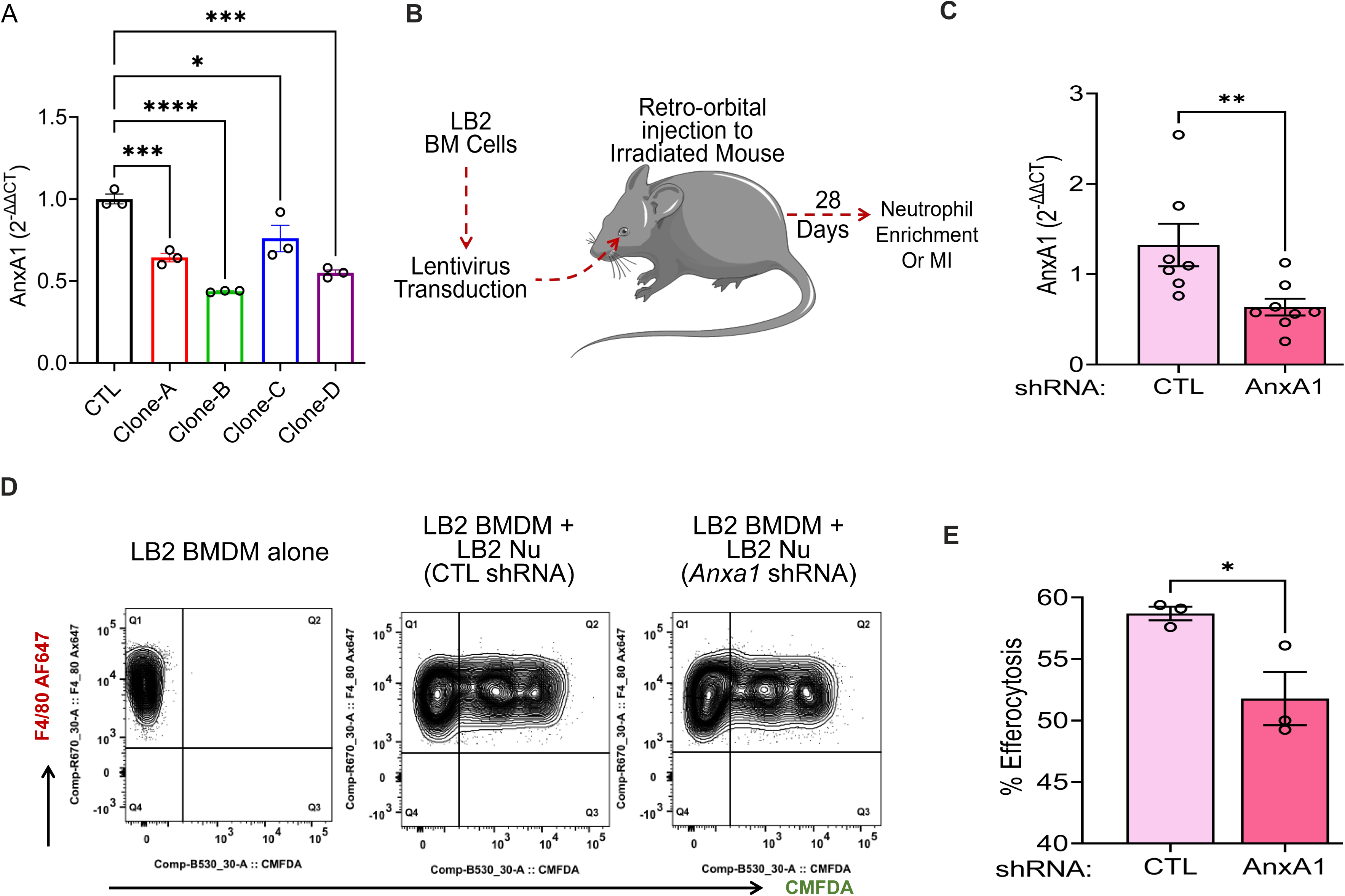
Lentivirus shRNA-mediated AnxA1 downregulation in neutrophils reduces their efferocytosis. (A) BMDM underwent lentivirus transduction of various clones (A-D) of AnxA1 shRNA versus control (CTL) shRNA for 48h, after which RT-qPCR was used to assess gene silencing efficiency, n=3/group. (B) Schematic depicting the experimental design to test the impact of AnxA1 knockdown in LB2 bone marrow-derived Nu. Briefly, LB2 BM was transduced with lentivirus containing either CTL shRNA or AnxA1 shRNA (1:1 mixture of clones B and D), after which they were retro-orbitally injected into lethally irradiated LB2 mice to generate bone marrow transplant (BMT) mice featuring LB2 bone marrow with or without AnxA1 knockdown, as confirmed via RT-qPCR analysis after terminal experiments (C), n=7 (CTL shRNA), n=8 (AnxA1 shRNA). 4 weeks after BM reconstitution, CTL shRNA and AnaA1 shRNA Nu were enriched from the respective BMT mice using magnetic columns, after which they underwent efferocytosis assays with LB2 BMDM (as described in Fig. 5) and flow cytometry analysis, as shown via contour plots (D) of LB2 BMDM incubated without Nu (left) or with CMFDA+ Nu from either CTL shRNA (middle) or Anxa1 shRNA (right) LB2 BMT mice. (E) Quantification of CTL shRNA versus Anxa1 shRNA LB2 Nu uptake efficiency into LB2 BMDM in terms of % efferocytosis, n=3/each group. Data are Mean ± SEM of independent experiments. ns, non-significant, * p < 0.05, ** p < 0.01, *** p < 0.001, **** p < 0.0001, One way ANOVA with Tukey’s post-hoc test (A), or Unpaired student’s t-test (C, E).

### Lentivirus-mediated AnxA1 deletion prevents the ameliorative effects of myeloid cell-specific β2AR deletion on post-MI outcomes in vivo

Since AnxA1 knockdown decreased the *in vitro* efferocytosis efficiency of LB2 myeloid cells, we hypothesized that the ameliorative post-MI effects observed in LB2 mice may be prevented via shRNA-mediated knockdown of *Anxa1*. Following the protocol described in Fig 6B, we generated chimeric LB2 mice with CTL versus Anxa1 shRNA-transduced BM. Four weeks following BMT, the mice underwent MI surgery and cardiac function and remodeling responses were assessed. Echocardiography showed a significant reduction in both %EF and %FS at 3- and 14-days post-MI in *Anxa1*-shRNA versus CTL-shRNA LB2 BMT mice (Fig. 7A, B). Additionally, LV wall thinning and chamber dilatation trended toward worsening in the *Anxa1*-shRNA versus CTL-shRNA LB2 BMT mice (Supplemental Fig 3). Moreover, MT staining revealed that the % infarct length as well as border zone interstitial fibrosis were both significantly increased in the *Anxa1*-shRNA versus CTL-shRNA LB2 BMT mice at 14 days post-MI (Fig 7C-F). Thus, knockdown of *Anxa1* in BM cells was sufficient to prevent the ameliorative effects of myeloid cell-specific β2AR deletion on cardiac function and fibrotic remodeling observed post-MI in LB2 mice (Fig. 1 and 2). In all, our data reveal a previously unrecognized role for β2AR in the regulation of myeloid cell-dependent efferocytosis in the heart following injury, wherein its deletion de-represses Anax1 expression to allow for enhanced efferocytosis-mediated Nu clearance to limit infarct expansion and loss of function (Fig. 7G).

**Figure 7.**
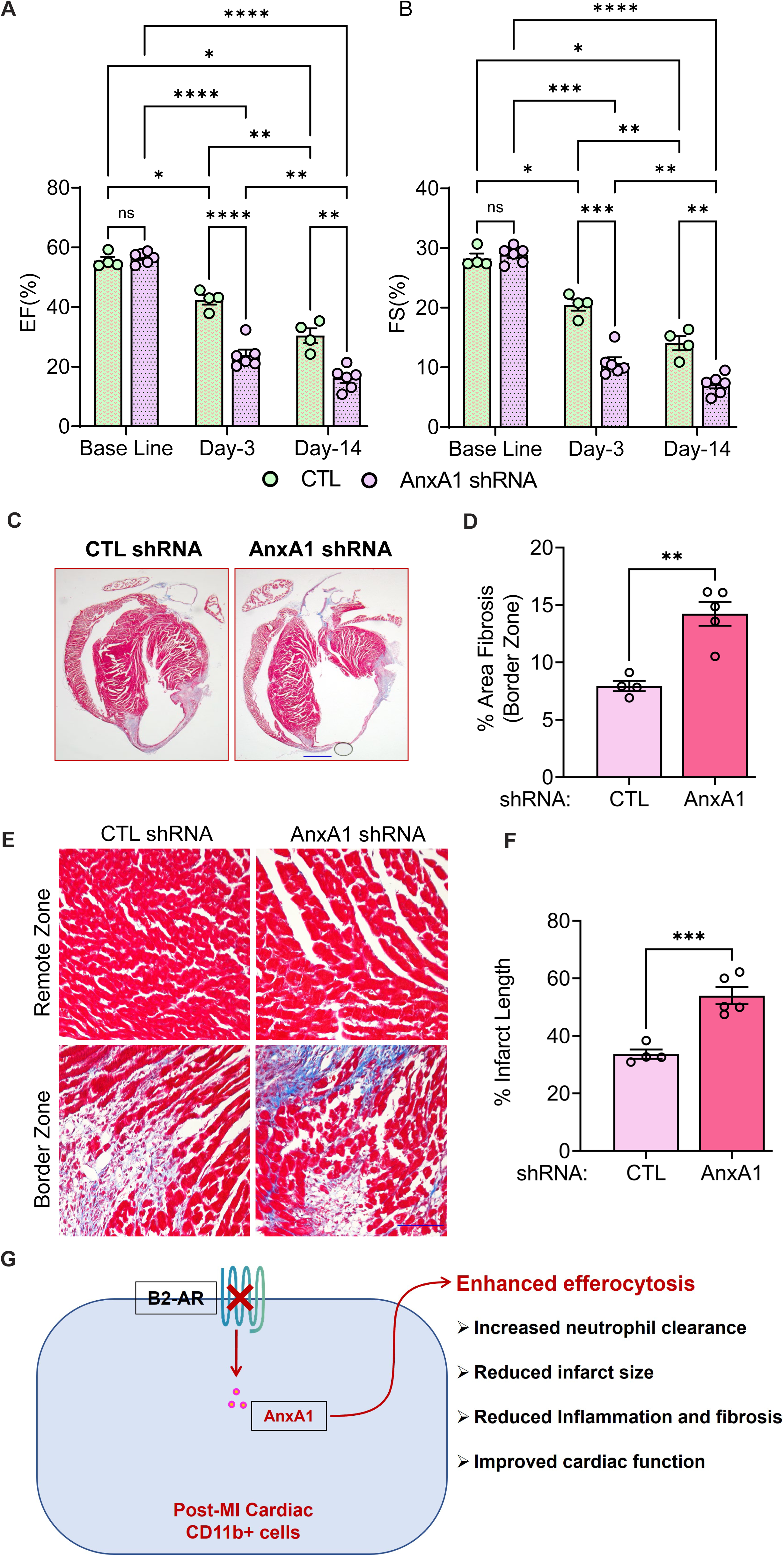
Lentivirus-mediated AnxA1 downregulation prevents the ameliorative effects of myeloid cell-specific β2AR deletion on post-MI outcomes in vivo. CTL shRNA and AnxA1 shRNA-transduced LB2 BMT mice were generated as described in Fig. 6, and, 4 weeks after BM reconstitution, underwent MI surgery. Histograms depicting %EF (A) and %FS (B) at baseline (BL), 3- and 14-days post-MI in CTL and AnxA1 shRNA LB2 BMT mice, n=4 for CTL, n=6 for AnxA1 shRNA. (C) MT staining of whole heart displaying infarct size of CTL and AnxA1 shRNA LB2 BMT mice at 14-days post-MI with quantification shown in (D), n=4 for CTL and n=5 for AnxA1 shRNA group, scale bar = 1000 µm. (E) Representative light microscopic images demonstrating border zone cardiac interstitial fibrosis in CTL and AnxA1 shRNA LB2 BMT mice at 14-days post-MI with quantification as % area fibrosis (F), n =4 for CTL and n=5 for AnxA1 shRNA group, scale bar = 100 µm. (G) Graphical abstract depicting working model of the cardioprotective impact of myeloid specific β2AR deletion, wherein loss of β2AR leads to enhanced expression of AnxA1 and subsequently increased efferocytosis of dead/dying Nu in the infarct area, leading to more efficient Nu clearance from the site of injury with corresponding improvements in cardiac function and remodeling. Data are Mean ± SEM of independent experiments. ns, non-significant, *p < 0.05; **p < 0.01, *** p < 0.001, **** p < 0.0001, Two-way repeated measures ANOVA with Tukey’s multiple comparisons test (A, B) or Unpaired student’s t-test (D, F).

## Discussion

At steady state, myeloid cells comprise a small portion of the total heart by volume^3^, but the release of cytokines and chemokines upon acute ischemic injury leads to a dramatic increase in the recruitment of peripheral myeloid cells, including Nu and Ly6C^hi^ inflammatory Mon^48^. Recruitment of these cells to the site of injury occurs quickly, within hours and persisting for several days, allowing the release of many factors including proteases and reactive oxygen species to digest and clear the infarct area^48, 49^, without which stabilization of the scar cannot occur and rupture ensues. We previously reported that deletion of β2AR in all cells of hematopoietic origin via β2AR KO bone marrow transplantation (BMT) resulted in generalized disruption of immune cell responsiveness to injury, including a decrease in Nu and Mon/Mac accumulation in the heart following MI-induced injury that correlated with impaired wound healing and ultimately cardiac rupture^31, 32^. A subsequent study further demonstrated that BMT mice lacking β2AR expression in all cells of hematopoietic origin also displayed impaired recruitment of immune cells, including Nu, Mon and Mac, into the heart in response to chronic catecholamine stimulation, resulting in protection from maladaptive remodeling^50^. Together, these studies demonstrated that immune cell-expressed β2AR modulates important facets of acute and chronic cardiac stress, with detrimental versus beneficial outcomes attained in a model-specific context. Further, they suggested that myeloid cell-specific β2AR may be particularly important in these outcomes due to the impact of generalized immune cell β2AR depletion on Nu, Mon and Mac responses to either acute cardiac injury or global catecholamine elevation. However, in contrast to mice with β2AR deletion in all cells of hematopoietic origin^31^, in our current study we found that mice with myeloid cell-specific β2AR deletion did not undergo cardiac rupture following MI and in fact displayed better contractile function than the control lines, more so early after injury, but also by 2 weeks post-MI, suggesting better scar stabilization and coinciding with decreases in fibrotic remodeling, infarct length and LV wall thinning.

Based on the previous reports discussed above^31, 32, 50^, it was expected that myeloid cell-specific β2AR deletion would result in their reduced recruitment to the heart after acute injury, however we did not observe a significant reduction in the accumulation of Nu, Mon or Mac in the heart within the first 24 hours post-MI. Thus, in contrast to the BMT model of β2AR deletion in all cells of hematopoietic origin, myeloid cell-specific β2AR deletion does not alter their responsiveness or ability to traffic to the ischemic heart immediately following injury. The reasons for this observation are unclear, but likely stem from the difference in models, wherein the adoptive transfer of hematopoietic stem cells lacking β2AR expression would result in the clonal expansion of all immune cell lineage precursors, except tissue resident macs. A previous microarray study that analyzed bulk bone marrow cells lacking both β1AR and β2AR reported thousands of differentially expressed genes involved in myriad ontological functions^51^. Thus, global deletion of β2AR in bone marrow stem cell precursors may result in a greater number and/or different alterations in myeloid cell transcriptomes, and ultimately responsiveness to injury, than lineage-specific deletion. Indeed, LysMCre mice were previously shown to induce reporter gene recombination in ∼40% splenic Macs, ∼40% peripheral Mon and ∼80% peripheral Nu versus 100% induction by Vav1-Cre that targets all hematopoietic cells, akin to the BMT model^52^, thereby providing a more selective gene deletion strategy with less severe effects that allows myeloid cell recruitment to the heart following injury.

Although ultimately required to prepare the injury zone for repair, a lack of removal of dead/dying Nu can prolong inflammation and aggravate further tissue destruction, thus efficient clearance of Nu is beneficial to transition the post-injury response away from the acute inflammatory phase toward repair and injury resolution^12, 49^. While myeloid cell-specific loss of β2AR did not alter the initial recruitment of Nu to the heart following injury, the accumulation of Nu was significantly reduced at 4 days post-MI, suggesting enhanced clearance. Indeed, expression of several efferocytosis-related genes were found to be increased in the bone marrow cells of mice with myeloid cell-specific β2AR deletion, including *Anxa1*, which was increased both at baseline and after MI. Additionally, *Anxa1* expression was increased in myeloid cells isolated from the hearts of mice with myeloid cell-specific β2AR deletion versus controls at 1-day post-MI, and Anxa1 protein was increased within individual myeloid cell types, including Nu, Ly6C^hi^ Mon and Mac, at 4 days post-MI. This increased expression across multiple myeloid cell types was notable since several studies in non-cardiac fields have demonstrated that myeloid cell (Nu or Mac)-produced Anxa1 acts to enhance Nu apoptosis and subsequent Mac-mediated efferocytosis to resolve inflammation^16–21, 45^, consistent with the enhanced Nu clearance observed at 4 days post-MI.

With respect to cardiac injury specifically, previous work has shown that ANXA1 is increased in human chronic heart failure patients^53^ and that global deletion of Anxa1 in mice worsens inflammation, fibrosis, angiogenesis and cardiac function following ischemic injury^54, 55^, whereas treatment of mice with recombinant Anxa1 or its N-terminal-derived peptide Ac-Anxa12-26 promotes cardiomyocyte survival, angiogenesis and better cardiac performance outcomes after ischemia^55–57^. These data are consistent with a pro-reparative role for Anxa1 via numerous mechanisms. Here, we found that in the context of acute ischemic cardiac injury, myeloid cell-mediated efferocytosis is also enhanced by Anxa1, with β2AR deletion causing higher myeloid cell-specific Anxa1 expression, increased efferocytosis *in vitro* and enhanced Nu clearance after cardiac injury *in vivo*, and knockdown of Anxa1 expression reversing these effects. Of note, it was previously suggested that Anxa1 may also act to reduce Nu extravasation, thereby dampening the inflammatory response after injury^58^, however in our study we did not observe a decrease in Nu accumulation in the heart following MI, suggesting that initial Nu extravasation from the peripheral circulation remains intact in mice with myeloid-specific β2AR deletion. However, due to the pleiotropic nature of Anxa1, as outlined above, it is possible that following the pro-efferocytosis effect we observed, additional beneficial effects of β2AR deletion-mediated enhancement of myeloid cell-specific Anxa1 expression could directly influence other remodeling processes such as fibrosis, which may warrant follow-up investigation.

Notably, while Anxa1 expression is known to be acutely induced by glucocorticoid (GC) stimulation, cAMP-elevating agents have also been demonstrated to increase Anxa1 expression in Nu to decrease their survival and enhance their clearance^59^. Despite both GC and cAMP-elevating agents being capable of increasing Anxa1 expression, it has been suggested that this may be mutually competitive at the promoter level^60^. In addition, the Anxa1 promotor itself has been suggested to contain binding sites for transcriptional repressors, such as PU.1, which can be de-repressed via interaction with other transcriptional mediators thereby allowing Anxa1 expression^61^. In our study we found that genetic deletion of myeloid cell-specific β2AR altogether resulted in enhanced Anxa1 expression, suggesting that β2AR likely acts to repress Anxa1 expression and that its absence relieves this negative regulation. While the intermediate regulator(s) of this effect remain to be determined, since cAMP-elevating agents promote Anxa1 expression, it is likely that other β2AR-sensitive signaling pathways mediate this basal repression. Another major consideration in the context of cardiac injury is the rapid and excessive release of local catecholamines in the inured myocardium where recruited myeloid cells accumulate^62^. In this scenario, high levels of catecholamines would be expected to quickly desensitize β2AR and/or promote a switch in its coupling from the cAMP-stimulating Gαs protein to the cAMP-inhibiting Gαi protein^63^, mechanisms that in their own right may normally act to dampen Anxa1 expression but that would be absent in the context of β2AR deletion, which will require further investigation.

While we observed that LysM-Cre-mediated deletion of β2AR resulted in enhanced Anxa1 expression in myeloid cells both before and after cardiac injury, resulting in increased efferocytosis of Nu in vitro and post-MI Nu clearance in vivo, a limitation of our study stems from the LB2 mouse model itself, in which β2AR deletion in all myeloid cells does not indicate whether its loss in a single myeloid cell subtype is sufficient to drive the phenotype. Lentivirus-mediated downregulation of Anxa1 in LB2 bone marrow prior to transplantation into LB2 mice was able to restore normal efferocytosis *in vitro* and post-MI fibrotic cardiac remodeling and dysfunction *in vivo*, suggesting that the peripherally recruited cells are responsible for the effect, but still does not provide resolution between Nu and Mon. Thus, a logical extension of this work would be the deletion of β2AR using additional models of Cre expression driven by other cell type-selective promoters, such as those more selective for Nu, Mon and/or Mac. Since Anxa1 has been shown to promote ameliorative effects on cardiac remodeling after injury that involve both Nu- and Mac-related processes^55–57^, such models would resolve whether the β2AR deletion-mediated effect on Anxa1 expression in either cell type would be sufficient to improve post-MI injury resolution, or if changes in myeloid cells in general are required for maximal benefit.

Altogether, myeloid cell-specific deletion of β2AR promotes improved outcomes following acute cardiac injury, wherein deletion of β2AR de-represses Anxa1 expression to enhance efferocytosis and clearance of Nu in the days following injury, thereby dampening cardiac dysfunction and fibrotic remodeling. Thus, we have revealed a previously unrecognized role for β2AR in the regulation of myeloid cell-dependent efferocytosis in the heart following injury, which may provide a potential new point of regulation of acute cardiac remodeling responses following ischemic injury.

## Funding

This work was supported by National Institutes of Health (R01 HL139522 and P01 HL147841 to DGT) and the American Heart Association (Postdoctoral Fellowships 909041 to TKN and 20POST35180177 to AB).

## Notes

### Competing Interest Statement

The authors have declared no competing interest.

