## Supplemental data for "Loss of Myeloid Cell-Specific β2-Adrenergic Receptor Expression Ameliorates Cardiac Function and Remodeling after Acute Ischemia"

**Supplemental Material**

**Supplementary Figure 1.** Graphs depicting various echocardiographic parameters (A; HR, B; SV, C; Co, D; LV mass, E; LVAWs, F; LVAWd, G; LVPWs, H; LVPWd, I; LV diameter-s, J; LV diameter-d, K; Volume-s, L; Volume-d) in sham and 3- and 14-day post-MI FL/FL, LMC and LB2 mice. Data are Mean ± SEM of independent experiments. ns, non-significant, $*p < 0.05$, Two-way ANOVA (mixed model) with Tukey's post-hoc test. n=11 for sham FL/FL, n=9 for sham LMC, n=12 for LB2, n=10 for MI FL/FL, n=5 for MI LMC, n=11 for MI LB2 (day 3), n=12 for MI FL/FL, n=9 for MI LMC, n=6 for MI LB2 (day 14).

**Supplementary Figure 2.** FACS pseudo color plot analysis showing the enrichment of bone marrow Nu using CD11b BV650 and Ly6G BV421 antibodies.

**Supplementary Figure 3**. Histogram depicting various parameters (A; HR, B; SV, C; Co, D; LV mass, E; LVAWs, F; LVAWd, G; LVPWs, H; LVPWd, I; LV diameters-s, J; LV diameters-d, K; Volumes-s, L; Volumes-d) of echocardiography reading in CTL and 3- and 14 days post-MI in LB2 BMT mice group. n=5 for CTL, n=6 for AnxA1 shRNA. Data are Mean ± SEM of independent experiments. ns, non-significant, $*p < 0.05$, One way ANOVA with Tukey's post-hoc test.

**Supplemental Figure 1**

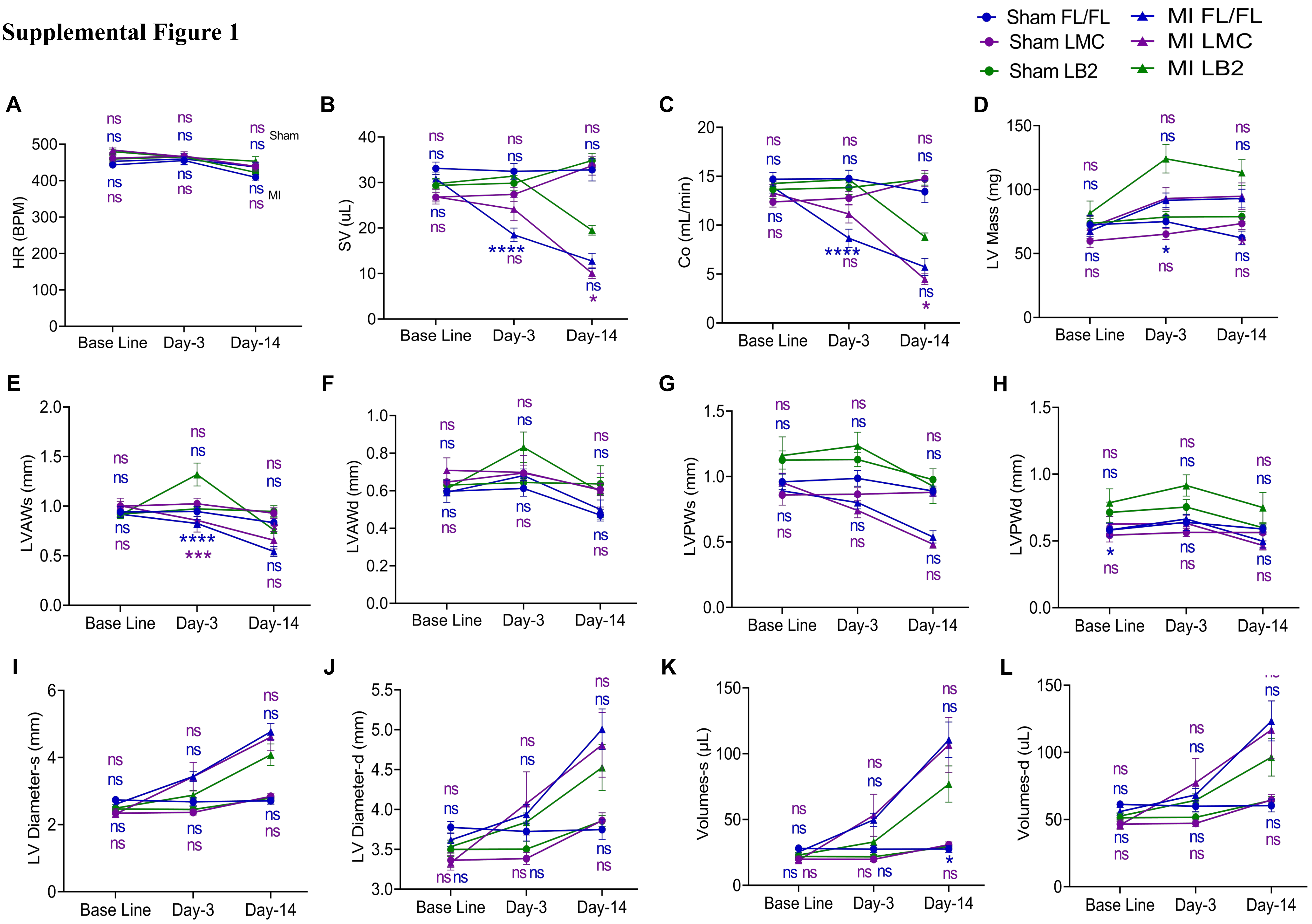

Supplemental Figure 2

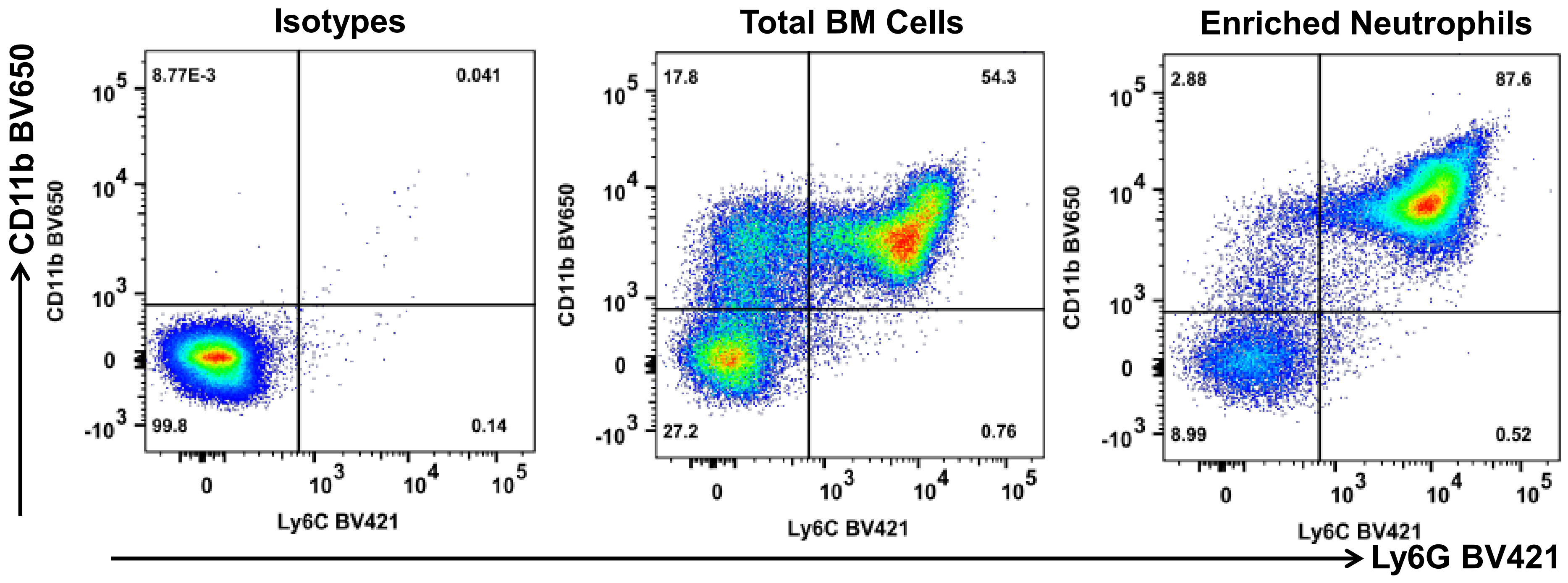

**Supplemental Figure 3**

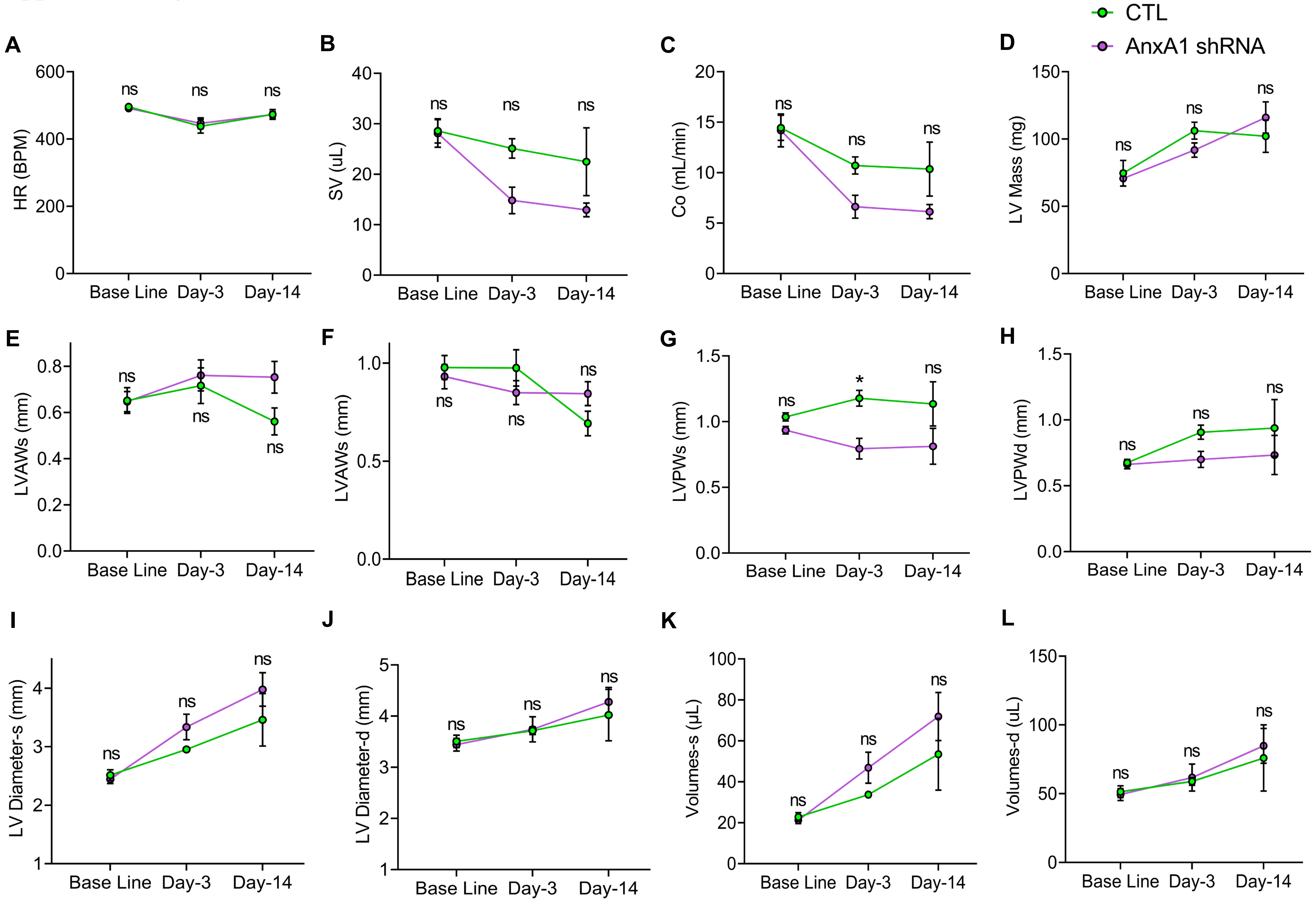

**Supplemental Table 1:** List of RT-qPCR primers.

| Gene Name | Primer Sequence (5’----------3’) |  |
|---|---|---|
| | **Forward** | **Reverse** |
| Flox | CCAAAGTTGTTGCACGTCAC | GCACACGCCAAGGAGATTAT |
| Cre | GGCGTTTTCTGAGCATACCT | CTACACCAGAGACGGAAATCCA |
| Gapdh | AACAGCAACTCCCACTCTTC | CCT GTT GCT GTA GCC GTA TT |
| Adrb2 | AAG AAT AAG GCC CGA GTG GT | GTA GGC CTG GTT CGT GAA GA |
| AnxA1 | TGTCAGGACTTGAGTGTGAA | GCTCCTACTGGTCAGAATTG |
| Mertk | CAGTCACACCTGAAAAGGAT | AAAAGTTCATCTGGTGATGG |
| Axl | GAGGAAGAAGGAGACTCGAT | TCCTTCAGCTCTTCACTGAT |
| Stab1 | GACGAGCTCACCTACAAGAC | GGCATTGATACTCAGAAAGG |
| Stab2 | GATGAGCCATACACCATTTT | TCTGACGAGTTCCAGAAGTT |

**Supplemental Table 2:** List of antibodies used for flow cytometry.

| Antibodies | Fluorophore | Manufacturer | Dilution |
|---|---|---|---|
| CD45 | BUV395 | BD, NJ, USA | 1:100 |
| CD11b | BV650 | BD, NJ, USA | 1:50 |
| Ly6C | PE-Cy7 | BioLegend, CA, USA | 1:300 |
| Ly6G | BV421 | BD, NJ, USA | 1:100 |
| F4/80 | Alexa Fluor 647 | Invitrogen, CA, USA | 1:100 |
| AnxA1 | PE | abcam, Cambridge, UK | 1:50 |
| Rat IgG2b | BUV395 k | BD, NJ, USA | 1:100 |
| Rat IgG2b | BV650 k | BD, NJ, USA | 1:100 |
| Rat IgG2c | PE-Cy7 k | BioLegend, CA, USA | 1:300 |
| Rat IgG2a | BV421 | BD, NJ, USA | 1:100 |
| Rat IgG2a | Alexa Fluor 647 k | Invitrogen, CA, USA | 1:50 |
| Rab mAb | PE | abcam, Cambridge, UK | 1:50 |

**Supplemental Table 3:** Details of Mouse AnxA1 shRNA RFP-Lentivirus plasmids target sequences with TR30030 (Origene, ) backbone with 5’TCAAGAG3’ as loop.

| AnxA1 shRNA Clone | Sequence |
|---|---|
| HC108601A | GGCAATGGTATCAGAATTCCTCAAGCAGG |
| HC108601B | GGCCAAAGACATCACTTCAGATACATCTG |
| HC108601C | TGACCGTTGTCAGGACTTGAGTGTGAATC |
| HC108601D | AAGCCATCCTGGATGAAACCAAAGGAGAC |

**Supplemental Table 4:** List of reagents, chemicals, and kits.

| Name | Catalog | Manufacturer |
|---|---|---|
| RPMI-1460 | 10-040-CV | Corning, NY, USA |
| DMEM | 10-117-CV | Corning, NY, USA |
| FBS | 900-108 | GeminiBio, CA, USA |
| PSF | 400-101 | GeminiBio, CA, USA |
| PBS | PB399-20 | Thermo Fisher Scientific, MA, USA |
| L-Ascorbic acid | A5960 | Sigma Aldrich, MO, USA |
| Isoproterenol | 16504 | Sigma Aldrich, MO, USA |
| Collagenase | L5004159001 | Worthington, NJ, USA |
| DNase-I | 10104159001 | Sigma Aldrich, MO, USA |
| Hyaluronidase | H3506 | Sigma Aldrich, MO, USA |
| EDTA | 351-027-101 | Quality Biologicals, MD, USA |
| ACK lysing buffer | 118-156-721 | Quality Biologicals, MD, USA |
| Live/Dead Aqua | L34966 | Thermo Fisher Scientific, MA, USA |
| PureLink™ RNA Mini Kit | 12183018A | Thermo Fisher Scientific, MA, USA |
| High-Capacity cDNA Reverse Transcription Kit | 4368813 | Thermo Fisher Scientific, MA, USA |
| PowerUp™ SYBR™ Green Master Mix for qPCR | A25741 | Thermo Fisher Scientific, MA, USA |
| CD11b Microbeads | 130-097-142 | Miltenyi Biotec, Bergisch Gladbach, Germany |
| Neutrophil Isolation Kit, mouse | 130-097-658 | Miltenyi Biotec, Bergisch Gladbach, Germany |

**Supplemental Table 5:** List of abbreviations.

| Abbreviations | Full Name |
|---|---|
| FBS | Feta Bovine Serum |
| DMEM | Dulbecco's Modified Eagle Medium |
| RPMI-1640 | Roswell Park Memorial Institute |
| EDTA | Ethylenediaminetetraacetic acid |
| PBS | Phosphate-buffered saline |
| BMDM | Bone Marrow Derived Macrophages |
| SFM | Serum Free Medium |
| ACK buffer | Ammonium–chloride–potassium lysing buffer |
| BL | Base Line |
| ISO | Isoproterenol |
| shRNA | Short hairpin RNA |
| RFP | Red Fluorescent Protein |
| cDNA | complementary Deoxyribonucleic Acid |
| RNA | Ribonucleic Acid |
| RT-qPCR | Real-time polymerase chain reaction |
| GAPDH | Glyceraldehyde 3-phosphate dehydrogenase |
| β2AR | β2-Adrenergic Receptor |
| BMT | Bone Marrow Transplant |
| ACS | Acute Coronary Syndrome |
| HF | Heart Failure |
| IACUC | Institutional Animal Care and use Committee |
| LCA | Left Coronary Artery |
| MT staining | Masson's Trichrome staining |
| DAMPs | Damage-Associated Molecular Patterns |
| TLR4 | Toll-like Receptor 4 |
| IL | Interleukin |
| CCR | CC Chemokine Receptor |
| CXCL | Chemokine (C-X-C motif) Ligand |
| CD | Cluster of Differentiation |
| M-CSF | Macrophage Colony-Stimulating Factor |
| IgG | Immunoglobulin G |
| BMP | Beats per minutes |
| SV | Stroke Volume |
| EF | Ejection Fraction |
| FS | Fractional Shortening |
| CO | Cardiac Output |
| LV | Left Ventricle |
| LVAW;s | Left ventricular anterior wall thickness; systole |
| LVAW;d | Left ventricular anterior wall thickness; diastole |
| LVPW;s | Left ventricular posterior wall thickness; systole |
| LVPW;d | Left ventricular posterior wall thickness; diastole |
| HR | Heart Rate |
| CTL | Control |
| SEM | Standard Error of the Mean |
| ANOVA | Analysis of Variance |

| | |
|---|---|
| HW | Heart Weight |
| TL | Tibia Length |
